# Leveraging AI for improved reproducibility of mathematical disease models: insights from a retinitis pigmentosa case study

**DOI:** 10.1101/2024.04.26.591317

**Authors:** Éléa Thibault Greugny, Nicolas Ratto, Loïc Etheve, Jean-Baptise Gourlet, Frédéric Cogny, Claudio Monteiro

## Abstract

Mathematical modeling of disease and drug action is becoming an indispensable component of drug development, underscored by recent examples of models predicting trial results. To be able to rely on such approaches, decision-makers need to be able to verify those results independently with in silico confirmatory studies. Artificial Intelligence (AI) offers a valuable avenue for improving the reproducibility of complex models, enabling their swift and software-agnostic deployment. This paper highlights AI’s impact through a case study on an Ordinary Differential Equation (ODE) model of Retinitis Pigmentosa (RP). We use a version of Chat GPT 4, a sophisticated large language model (LLM) developed by OpenAI, as customized by Mathpix company with additional capabilities. This setup facilitated the extraction of equations from the PDF and converted into a human-readable, text-based definition language called Antimony, which is part of the Python package tellurium. Subsequently, the model was converted into Systems Biology Markup Language (SBML) using tellurium and uploaded onto the jinkō platform for simulation. The RP model was efficiently and accurately implemented using AI techniques. Furthermore, we were able to reproduce the model behavior presented in the literature. Our findings advocate for the broader application of AI in mathematical model re-implementations to ensure reliability and reproducibility of the results.

## Introduction

The issue of irreproducible models has been a recurrent concern for the last decades (1, 2). A recent example in biology comes from the “Reproducibility Project: Cancer Biology” (3), which focuses on replicating experimental results from several high-profile cancer biology papers. As Errington et al. (4) articulate, scientific advancement relies not only on innovation but also on the verification of results: “Innovation without verification is likely to accumulate incredible results at the expense of credible ones and create friction in the creation of knowledge, solutions, and treatments.” Beyond the costs of innovation, significant expenses are also associated with the verification stages (5). Holding many promises, especially in the causal consideration of complex phenomena, the field of Quantitative Systems Pharmacology (QSP) shares these same reproducibility challenges (6). However, with the recent widespread adoption of artificial intelligence (AI), there is potential for significant reductions in both cost and time required to reproduce results from QSP models documented in the literature. In the ever-evolving landscape of AI, the capability to swiftly and accurately transform theoretical models from research papers into functional algorithms is critical. The diversity in programming languages and formats complicates the verification of model results, even when code is provided. Moreover, having both the code and the equations does not ensure that the implementation accurately reflects the authors’ intentions as described in the research. This paper delves into the innovative domain of automatic model implementation using AI, a field poised to drastically accelerate research and its applications across various scientific and technological areas. By leveraging advanced machine learning techniques and natural language processing capabilities, AI systems can interpret, extract, and implement complex concepts and methodologies directly from scientific articles. This approach not only democratizes access to cutting-edge research by mitigating the need for deep technical expertise but also underscores the potential of AI in bridging the gap between theoretical research and practical application.

### Implementing model is long and difficult

Implementing QSP models from published papers into practical applications represents a unique set of challenges, primarily due to the completeness and inaccuracy of the information presented in the papers. Often, papers may outline the theoretical framework or model but omit fundamental details such as specific equations, initial conditions, or parameters, leading to significant discrepancies between the intended and the implemented models (6). Researchers from the European Molecular Biology Laboratory European Bioinformatics Institute (EMBL-EBI) (7), identified missing parameter values and initial conditions as the primary obstacle to reproducing published results.

According to Tiwari et al., among the 49% of models that were not directly reproducible, 9% could eventually be reproduced after manually correcting errors found in the manuscripts, such as typographical errors in parameters values and units or missing terms (7). This necessitates not only the identification of potential errors but also a profound understanding of the model to correct them.

These challenges highlight the importance of meticulous scrutiny and a deep understanding of the underlying principles when attempting to implement models from academic papers. They also underscore the potential value of tools and methodologies that could assist in accurately interpreting and implementing these models, thereby narrowing the gap between theoretical research and practical application.

Even when equations and parameters are extensively detailed in papers, the manual implementation of these models into a working system is both time-consuming and prone to human error. Each step, from coding the equations to establishing the initial conditions, requires precision and a deep understanding of both the theoretical model and the practical computing environment. The time investment for such an endeavor is substantial and susceptible to the errors previously mentioned.

To address the issue of manually re-coding models from equations in manuscripts, researchers advocate two main solutions: sharing model code in public repositories and publishing models in standard formats like SBML or CellML (8). These strategies are encapsulated in questions 4, 5, and 6 of the reproducibility scorecard designed by Tiwari et al. Nonetheless, among 110 sampled models, less than 30% had their mathematical expressions and simulation code made publicly available, and only 17% provided the model in a standard format (7). Moreover, a 2018 study by Stodden et al. revealed that merely providing materials upon request is inadequate: among 204 papers published in Science, artifacts were obtained from only 44%, with reproducibility confirmed for only 26% (9).

Still, these solutions are not perfect: public code may require proprietary, non-free software (e.g., MATLAB), and a format like SBML may not capture all the specificities of the model (e.g., solving details). Overcoming these issues requires additional information, which can be laborious work.

This reality emphasizes the challenges in bridging the gap between theoretical models in academic papers and their practical, real-world applications. It underscores the need for more automated, error-resistant methods to translate academic research into actionable technological solutions.

### Implementation of a model using AI

Recent advancements in machine learning and natural language processing offer promising methods to facilitate model implementation from papers, especially to automate the extraction and translation of published equations into reusable code. Using AI can help to both accelerate model implementation and reduce the likelihood of manual transcription errors.

We demonstrate the practical application of such approaches through the implementation of a mathematical model of RP (10). RP is a hereditary eye condition characterized by the gradual loss of vision due to the progressive degeneration of rod and cone photoreceptor cells. It affects roughly 1 in 4000 individuals worldwide (11). The model in question is based on four differential equations with forteen parameters, describing the evolution of the number of normally functioning rods (Rn), non-functioning rods (Rs), cones (C), and the nutrient pool (T) they depend on. This method can be adapted to any model with comparable level of detail, regardless of its size. The initial step of model re-implementation involves extracting the mathematical equations and parameter values from the original paper. For this task, we used GPT Mathpix (12), available through GPT Plus or Team subscription on the GPT-store. GPT Mathpix is a customized version of Chat-GPT 4, a sophisticated large language model (LLM) developed by OpenAI, enhanced with a specialized knowledge base and additional capabilities through API calls to Mathpix (13). The Mathpix chatbot uses its natural language capabilities to convert user prompts into the right API calls leveraging mathpix optical character recognition (OCR) capabilities to convert images of equations or tables into LaTeX format. Therefore, the extraction step consists in prompting ‘Can you extract the 4 ODEs in LateX from this picture?’—with the image file containing the ordinary differential equations (ODEs).

Figure 1 illustrates the prompt and the chatbot’s response with the attached ODEs. Requesting a specific output format is a recommended practice when prompting a LLM, as is thoroughly reviewing the answer. Subsequently, the equations and parameters were converted into Antimony (14) using the same Chat-GPT 4 (version available in April 2024) conversation. Antimony is a human-readable domain specific language (DSL) used for model definition within the Python package tellurium. The readable nature of Antimony facilitates review by researchers compared to SBML (15). Meticulous inspection is required to prevent the introduction of errors into the translated model.

**Fig. 1.**
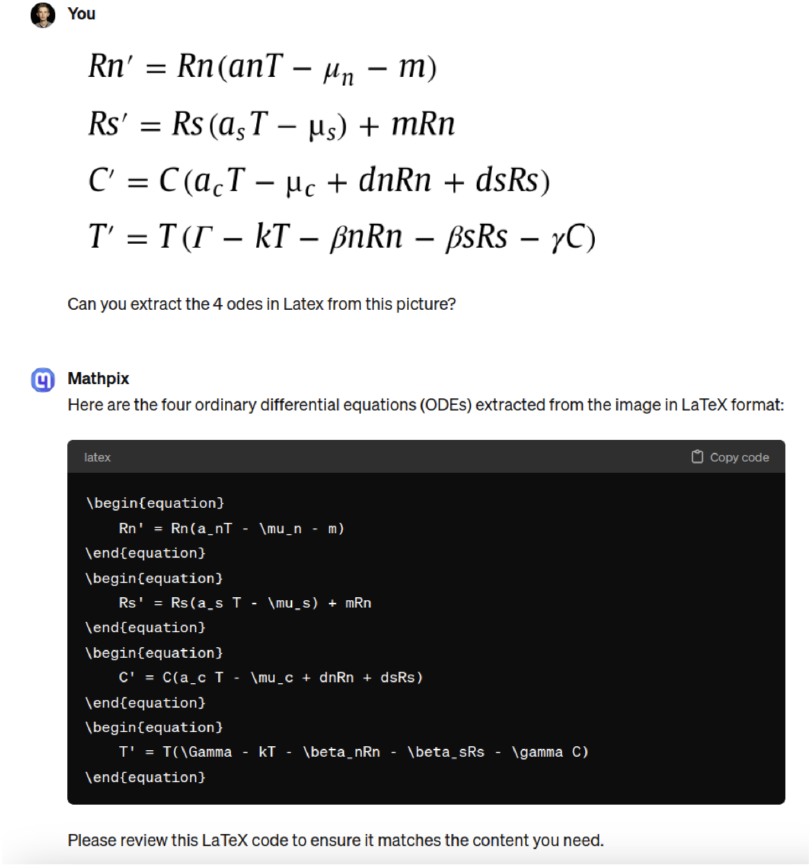
Screenshot of the conversation with GPT Mathpix to extract RP model equations in LATEX.

In Figure 2, Chat-GPT is asked to create the skeleton of the Antimony model from the previously extracted equations.

**Fig. 2.**
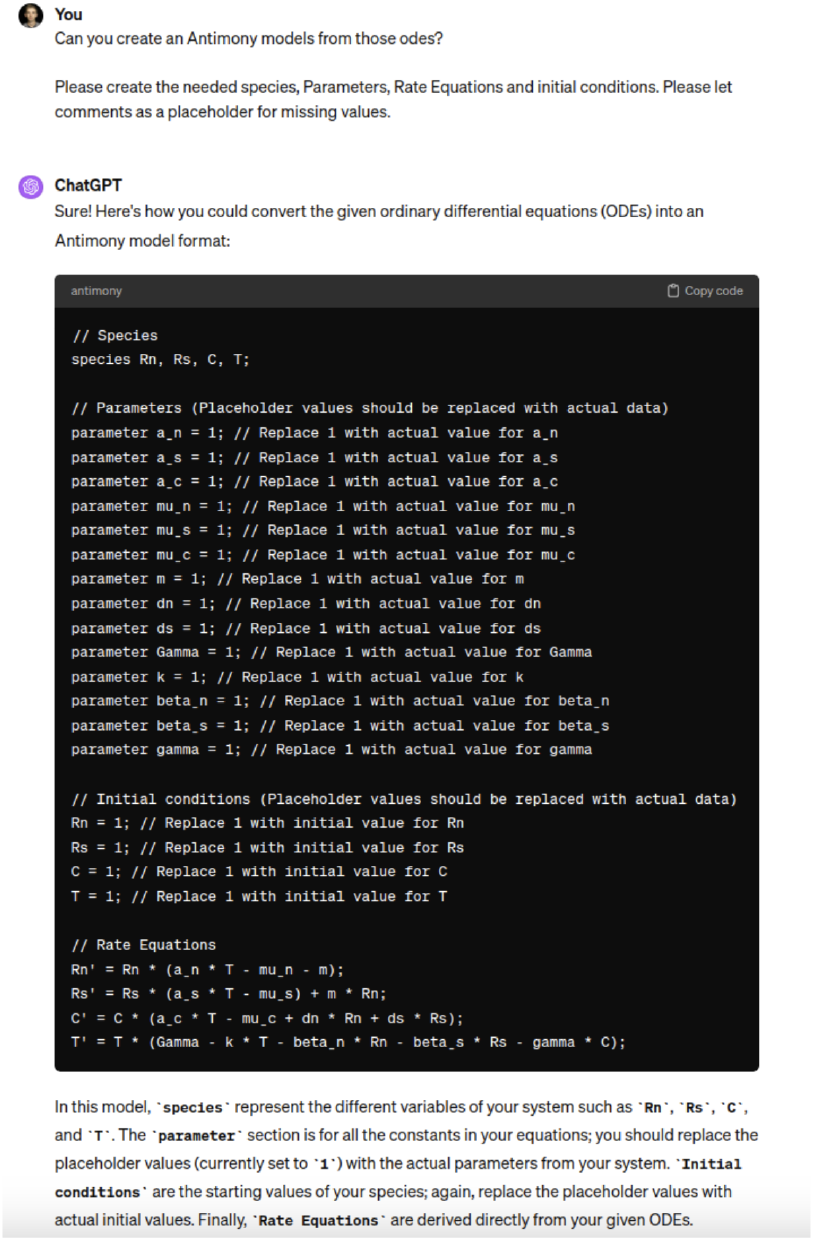
Screenshot of the ChatGPT-4 conversation to build an Antimony model from the extracted LaTeX equations.

For the RP model, the parameter values were extracted from the legend of Figure 6, and provided to Chat-GPT as a raw table (Figure 3). During this process, ChatGPT is able to detect missing parameters (parameter m) and initial conditions (16). After that, the model is created (Figure 3).

**Fig. 3.**
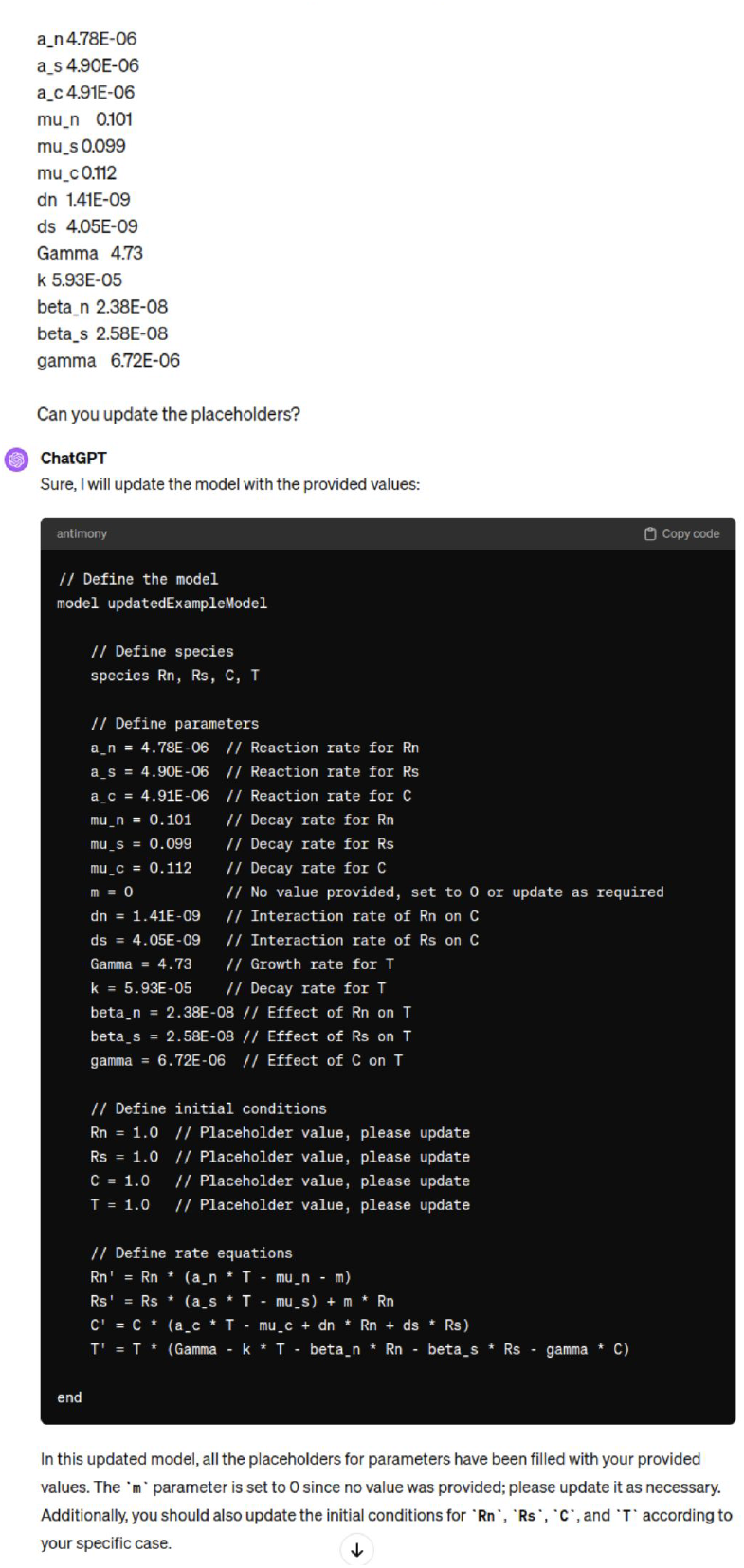
Screenshot of ChatGPT-4 conversation to specify parameter values and identify missing ones.

**Fig. 4.**
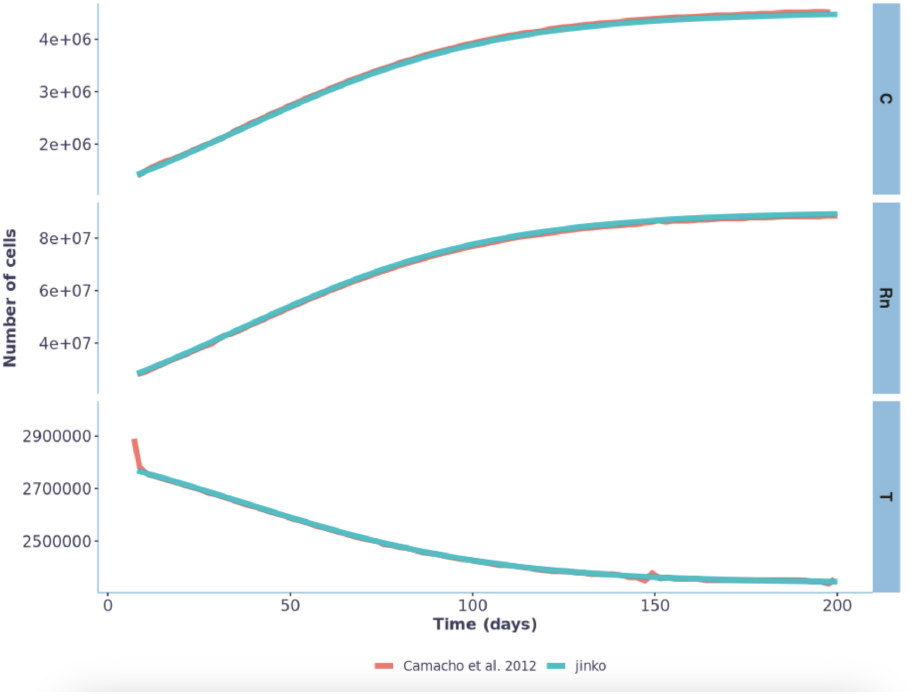
Recreating simulation outcomes originally presented by Camacho et al. (2013) using jinkō. The simulation results and parameter values were derived from Fig. 1 of Camacho et al. (16). Initial conditions have to be inferred from the graphs for simulation with jinkō, leading to slight variations.

After specifying the missing parameter values, the model written in Antimony was easily translated into SBML using tellurium (17, 18).

Translating a model to SBML holds significant interest to reuse published models. SBML’s standardized format ensures compatibility across various software platforms, allowing researchers to seamlessly integrate models into their preferred simulation and analysis tools. Here, the SBML model was uploaded on the jinkō platform, where it could be executed and further explored. Finally, the last but crucial step involves verifying that the simulated results reproduce the figure from the original paper. Here, the C, T, and Rn time series, as well as steady-states for the healthy eye, as depicted in Fig. 1 from Camacho et al. (16), were successfully reproduced (Figure 4).

### Known limits and attention points

Researchers can leverage AI to streamline the process of assimilating complex mathematical models, while remaining cognizant of the inherent limitations and critical aspects associated with this approach. Although a general method is outlined here, adjustments are often required to address the unique characteristics of specific cases.

It is noteworthy that not every paper includes units for all parameters and variables. Additionally, only a limited number of simulation solutions incorporate a unit-check or a unit conversion feature. These functionalities are essential to prevent the incorporation of incompatible units within equations, mitigating issues related to dimension errors or implicit conversion factors. These considerations become particularly crucial when reusing and adapting existing models.

One of the major challenges in leveraging IA to implement models from papers lies in the difficulty to retrieve missing information. Occasionally, a former paper from the authors is sufficient (like in RP) but often, reaching the authors remains necessary (7).

### Bridging Human Expertise and AI: Towards a RESTful API for Iterative QSP Model Development

LLMs have demonstrated remarkable proficiency in three key areas: 1) interpreting natural language, thereby offering an intuitive interface accessible to individuals without programming expertise; 2) converting natural language instructions into programmatic commands to utilize specialized, domain-specific applications, as demonstrated by the Mathpix GPT; and 3) maintaining a comprehensive conversational context, which supports a human-supervised, iterative development process. However, these models often struggle to generate lengthy outputs, a limitation particularly noticeable in the construction of extensive QSP models. This challenge is further compounded by the inherent randomness in the output of these generators.

Given these considerations, we propose that a significant advancement in the creation of step-by-step, reliable models from publications would require the development of a specialized programmatic API. This API would not only streamline the construction of models in a single session but also support iterative, granular modifications, thereby accommodating the necessity for incremental adjustments. Ideally, for seamless integration with the existing LLM infrastructure, such an API would be RESTful, featured with an OpenAPI specification (19), and complemented by a graphical user interface. This interface would provide users with immediate visual feedback on the execution of their commands. To our knowledge, no existing API fully meets these specifications. Nonetheless, we believe that merging natural language processing capabilities with solutions tailored for domain experts represents a logical and innovative direction for utilizing AI to iteratively develop dependable QSP models.

## Conclusion

In conclusion, the implementation of the RP model in this paper has demonstrated the remarkable capabilities of AI in translating complex equations into functional code, showcasing the practical potential of this approach in biomedical research.

However, our experience also underlines the inherent limitations posed by the quality of the input equations that can significantly impact the efficacy and reliability of the automated code generation process, highlighting the critical need for careful and comprehensive detailing in the theoretical frameworks provided in academic papers.

In addressing these challenges, it is important to recognize that formats capable of storing units, like the SBML, are already available. Furthermore, sophisticated software solutions, such as SimBiology in MATLAB or jinkō, offer advanced capabilities for automatically converting and managing these units during computations. The use of such tools and formats can significantly improve the accuracy and reusability of models derived from academic research.

While AI demonstrates remarkable efficiency in certain tasks, there are areas where its performance is lacking, necessitating manual intervention. As researchers striving to leverage AI productively, the full potential of AI lies in constructing a pipeline to enhance interactions between AI and human experts, rather than in attempting to force AI to integrate into existing human workflows. By crafting workflows that capitalize on the strengths of humans we can optimize productivity and innovation while mitigating the limitations inherent in each approach.

## ACKNOWLEDGEMENTS

This work was supported by Novadiscovery. All the authors of this article are employees of Novadiscovery. These authors contributed equally: Éléa Thibault Greugny and Nicolas Ratto.

